## Supplementary figures and images for "Geometry-Induced Competitive Release in a Meta-Population Model of Range Expansions in Disordered Environments"

### Supplemental Movie S1

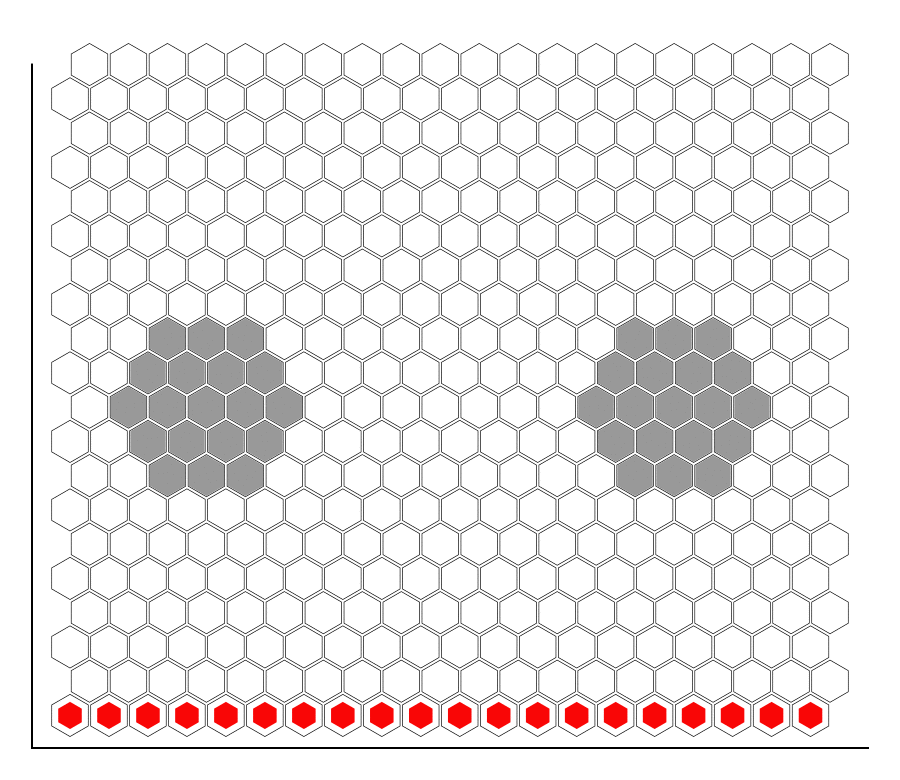

### Supplemental Movie S2

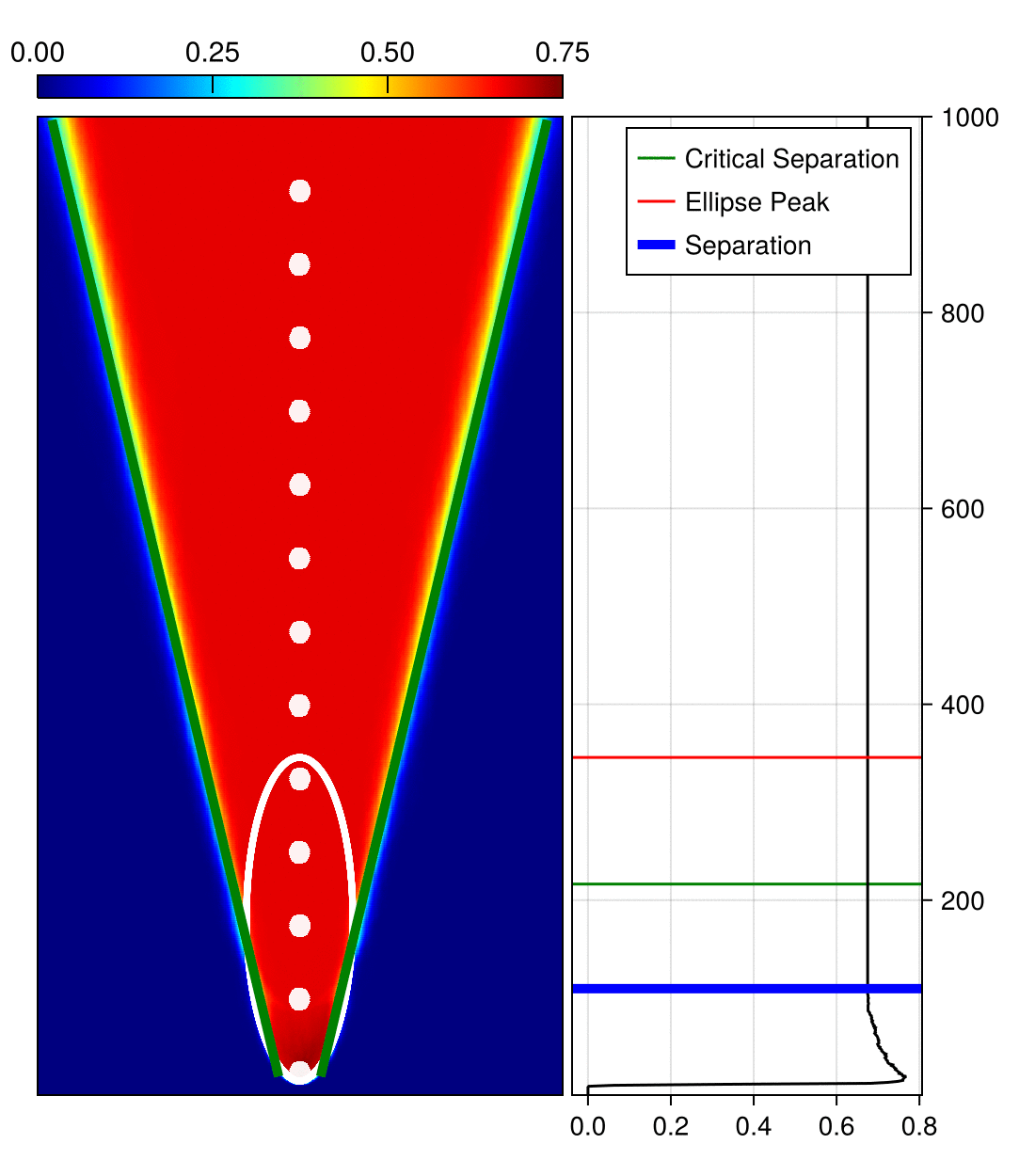
